# Lineage-specific CK2α deletion reshapes the transcriptome of hematopoietic stem cells toward an immune-primed state

**DOI:** 10.64898/2026.04.10.717787

**Authors:** Hannah Valensi, Rajesh Rajaiah, Marudhu Shanmugam, Daniyal Muhammad, Upendar Golla, Katherine Mercer, Anush Karampuri, Sinisa Dovat, Chandrika Behura, Yasin Uzun

## Abstract

Casein Kinase 2 (CK2) is a constitutively active kinase regulating proliferation and immune signaling and is frequently dysregulated in cancer, including acute myeloid leukemia (AML), making it a therapeutic target. CK2 comprises two catalytic subunits, CK2α or CK2α’, with two regulatory β subunits. The role of CK2α, the predominant catalytic subunit and principal mediator of CK2 kinase activity in hematopoietic cells, in steady-state hematopoiesis remains undefined. To define how CK2α shapes hematopoietic cells, we used bone marrow and spleen tissue samples of wild type control and conditional knock out (KO) of CK2α (*Csnk2a1*) in the hematopoietic compartment of transgenic mice. Using single-cell RNA sequencing, we profiled the transcriptomic changes associated with CK2α loss. Although HSC abundance was comparable between the control and CK2α-deficient samples, HSCs experienced the largest transcriptional response to CK2α loss among all cell types. CK2α-deficient HSCs displayed transcriptional remodeling for inflammatory and immune-associated programs, interferon signaling, and antigen presentation. Expression of inflammatory genes such as *S100a8* and *S100a9*, changed in opposite directions in bone marrow and spleen HSCs, demonstrating the transcriptional consequences of CK2α loss shaped by tissue context. Using a network-based approach, we identified immune-associated transcription factors *Nfkb1, Rfx5, Hes1*, and AP-1 family members as regulatory hubs driving these inflammatory transcriptional states in CK2α-deficient HSCs. Cell-cell communication profiling revealed multiple gains and losses in ligand-receptor communication between the HSCs and their immune microenvironment in KO. Our findings identify CK2α as a regulator of immune transcriptional programs in HSCs and suggest that disruption of CK2 signaling influences stem cell behavior and immune activation in contexts relevant to hematologic malignancies and CK2-targeted cancer therapies.

**Statement of significance:** This study reveals that inhibiting the protein CK2α forces blood stem cells into a stressed, immune-primed state. These tissue-specific findings highlight potential side effects for cancer therapies targeting this essential regulatory kinase.

## Introduction

Acute myeloid leukemia (AML) is an aggressive hematological malignancy characterized by impaired differentiation and uncontrolled expansion of myeloid progenitors (1). Despite advances in risk stratification and targeted therapies, long-term survival remains limited for many patients, particularly those with adverse cytogenetics or relapsed disease (2). AML arises through the accumulation of genetic and epigenetic alterations that disrupt normal lineage commitment and self-renewal programs, frequently involving pathways that regulate transcription, cell survival, and stress response (3,4). Increasing evidence supports the view that leukemic transformation co-opts molecular programs normally used by hematopoietic stem and progenitor cells (HSPCs), linking malignant progression to dysregulation of stem cell regulatory networks (5).

Hematopoietic stem cells (HSCs) reside at the apex of the hematopoietic hierarchy and sustain lifelong blood and immune cell production through tightly controlled decisions between quiescence, self-renewal, and lineage commitment (6). These decisions are governed by coordinated transcriptional and signaling programs that balance stability with responsiveness to environmental cues (7). In addition to maintaining hematopoietic output, HSCs actively participate in immune and inflammatory signaling, adopting transcriptional states associated with interferon responses and stress adaptation under both physiological and pathological conditions (8). Perturbation of these regulatory programs can bias lineage output and promote aberrant self-renewal, creating a permissive state for leukemic transformation (9).

Casein Kinase 2 (CK2) is a constitutively active serine/threonine kinase that contributes to a wide range of cellular functions. Rather than responding to acute signaling cues, CK2 maintains steady activity and plays roles in pathways that regulate transcription (10), protein synthesis (11), cell survival (12), and stress adaptation (13). Its catalytic subunits (CK2α or CK2α’) and regulatory subunits (CK2β) can operate within the canonical heterotetramer (αα-ββ or α’α’-ββ) or play independent roles, enabling CK2 to regulate an unusually broad set of substrates across many tissues (14). Because CK2 intersects with signaling networks frequently altered in cancer, inflammatory disease, and immune dysregulation, it has become an important molecular target in both basic biology and translational research (15). Notably, recent large-scale perturbation analyses have identified the CK2 inhibitor CX-4945 as a modulator of immune signaling, including upregulation of major histocompatibility complex class I (MHC-I) gene expression in immune-active contexts (16), suggesting a potential role for CK2 in regulating antigen presentation programs. Many transcription factors and signaling components that influence HSC fate decisions are subject to CK2-dependent regulation (17), raising the possibility that CK2 activity contributes to the balance between stem cell maintenance and lineage commitment.

Consistent with this role, CK2 has been implicated in leukemia pathogenesis and the maintenance of malignant stem cell populations (18). Pharmacologic inhibition of CK2 has demonstrated anti-leukemic activity in preclinical models and is currently under clinical investigation as a therapeutic strategy (19,20). While these findings support the consensus that CK2 is a candidate oncogenic dependency in AML (21), they also raise questions about how systemic CK2 inhibition may affect normal hematopoietic stem cell compartments. Prior studies using hematopoietic-specific CK2α deletion have shown that CK2α-deficient mice remain viable into adulthood and exhibit splenomegaly and expansion of HSC populations (17,22,23), in contrast to deletion of the CK2β subunit, which causes embryonic lethality and severe anemia due to disruption of GATA1 stability (24).

Single-cell RNA sequencing provides a powerful approach to resolve how genetic perturbations affect heterogeneous tissues by enabling simultaneous assessment of cell-type composition and transcriptional state (25). This approach is particularly well-suited for interrogating hematopoietic systems, where rare stem and progenitor populations coexist with diverse mature lineages (26). It also enables the construction of gene networks for these populations without the necessity of using multiple samples, to reveal the effect of perturbations in transcriptional signaling (27).

In this study, we performed hematopoietic-specific CK2α conditional knockout in Vav-iCre *Csnk2a1*^f/f^ mice. We focused particularly on HSCs because CK2 has been implicated in regulating stem cell survival, quiescence, and signaling pathways that are frequently dysregulated in AML (28). As the long-lived cells that sustain hematopoiesis and can serve as the cell of origin for leukemia (29), HSCs provide a relevant context in which to investigate the transcriptional and regulatory consequences of CK2α deficiency. To profile the transcriptome of hematopoietic cells, we obtained tissue samples from bone marrow and spleen, where these cell populations are densely populated. Particularly, bone marrow is the primary site of HSC residence and is closely linked to the initiation and progression of AML, whereas the spleen can function as a secondary hematopoietic organ and reservoir for immune and progenitor cells. Although both tissues harbor hematopoietic populations, they represent distinct microenvironments with differing oxygen tension, vascular organization, and immune exposure, raising the possibility that loss of CK2α elicits tissue-specific transcriptional responses.

## Results

To assess the downstream effects of CK2α gene perturbation in hematopoietic cells, we used samples from wild type control and hematopoietic-specific CK2α conditional knockout (KO) mice (**Fig. S1, S2**) bone marrow and spleen tissues and performed single-cell RNA sequencing for all samples (**Fig. 1A**). Using this novel dataset, we assessed the relative changes in cell populations in addition to transcriptional responses to KO, both in terms of individual genes and pathways. We also characterized differential gene expression, pathway activity, gene regulatory network (GRN) remodeling, and intercellular communication in HSCs following CK2α deletion (**Fig. 1B**).

**Figure 1.**
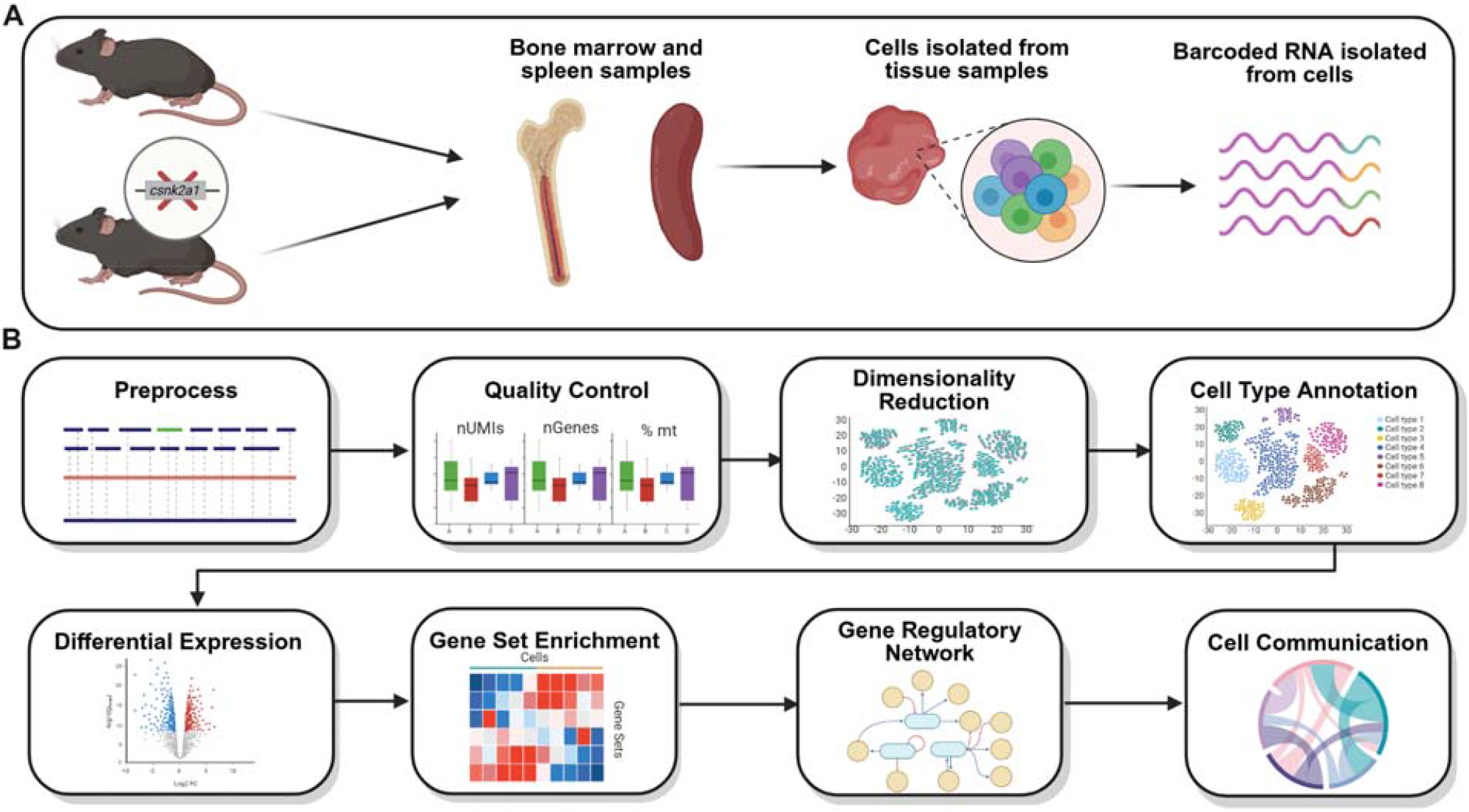
Workflow of the study. (**A**) Overview of the experimental workflow for processing the samples and sequencing. (**B**) Overview of the analytical steps for scRNA-Seq data.

### CK2α deletion maintains HSC abundance but promotes myeloid skewing

For the available tissue samples, we performed single-cell RNA sequencing (scRNA-seq) on BM and SPL tissues of WT control and KO mice. We removed the low-quality cells with strict quality-control metrics and downsampled each sample to an equal number of high-quality cells to avoid sample size bias in downstream analysis (**Fig. S3A-B**) ending with a large number of (7,025) cells for each individual sample.

We processed WT control and KO sample data for each tissue were processed together in an integrated manner using Seurat (30) (See Methods). We obtained 11 clusters in bone marrow and 12 clusters in spleen. Clusters were annotated to their respective cell types using expression of the canonical gene markers (**Fig. 2B**) in conjunction with differential gene expression across clusters queried against the Haemopedia database (31). We found dendritic, hematopoietic stem cells (HSC), macrophages, neutrophils, and plasma cells in both tissues, whereas mast cells were found in the bone marrow alone. Different subsets of specialized T cells, B cells and neutrophils were found in the two tissues. The number of genes expressed per cell type reflected the expected transcriptional diversity across hematopoietic populations (**Fig. S4**). Dimensionality reduction and clustering revealed distinct transcriptional states corresponding to known cell types (**Fig. 2A**). Control and CK2α knockout cells largely overlapped in UMAP space, indicating preservation of major lineages while allowing assessment of genotype-specific shifts in population composition (**Fig. 2C**).

**Figure 2.**
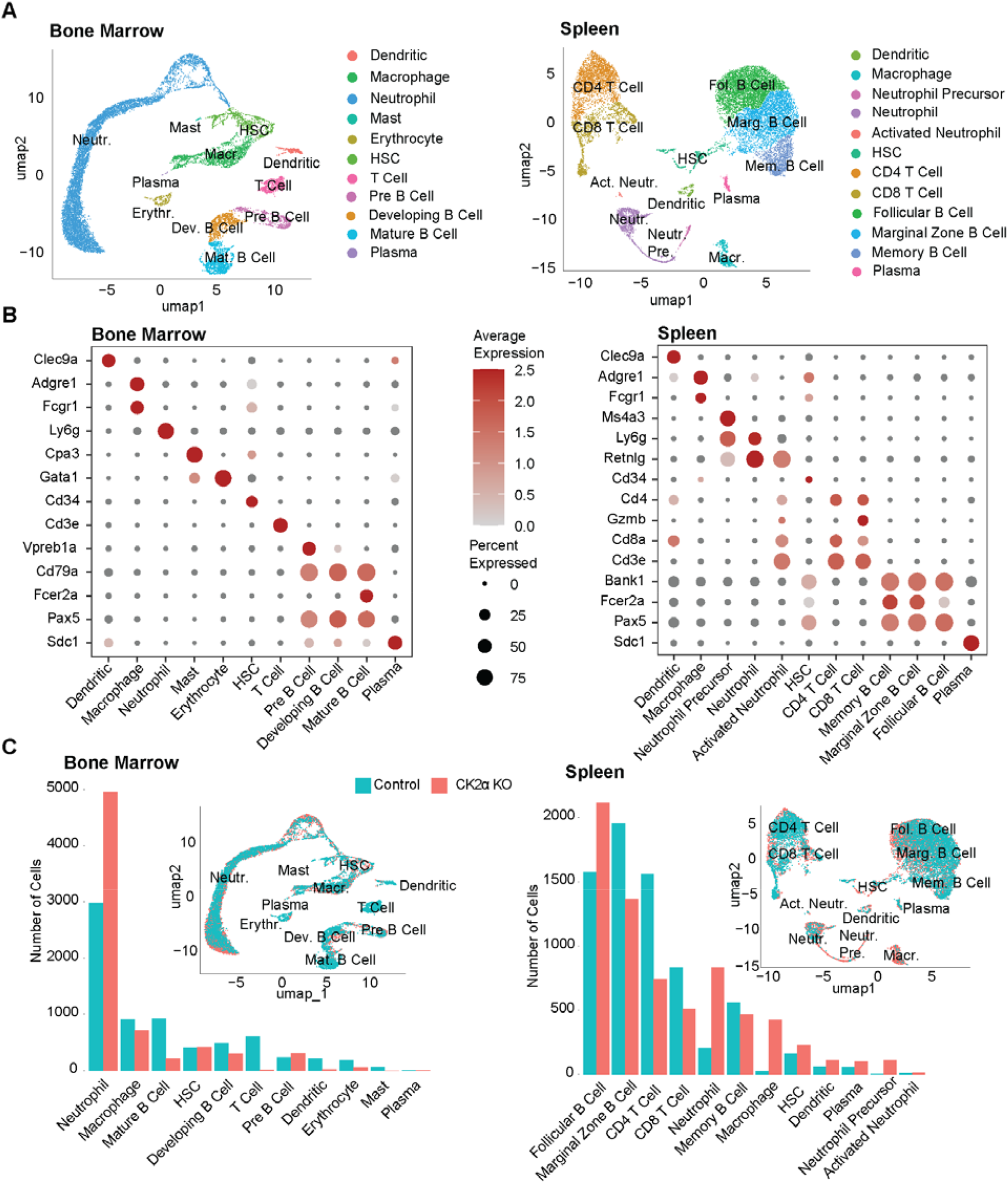
Cell clustering and cell type annotation. (**A**) UMAPs embeddings of hematopoietic precursors and immune cells from bone marrow and spleen, showing transcriptionally distinct clusters labeled by assigned cell type. (**B**) Dot plot of canonical marker genes used to validate cell type annotations across clusters for each tissue. Dot size represents the proportion of cells expressing each gene, and color intensity represents scaled expression. (**C**) Bar plots showing the number of cells in each annotated cell type, separated by control and knockout conditions for each tissue, and UMAPs for each tissue overlaid by condition of control or knockout.

We observed that cell-type abundances in control samples matched the expected hematopoietic composition (32). In bone marrow, neutrophils represented the dominant population, followed by macrophages and multiple B cell subsets, while T cells comprised a smaller fraction along with HSCs. In contrast, the spleen was enriched for lymphoid populations, with B cell subsets comprising the majority of cells and substantial contributions from CD4 and CD8 T cells. Neutrophils were comparatively low in terms of population in the spleen, consistent with its lymphoid-based composition.

CK2α deletion altered the relative abundance of specific hematopoietic populations without disrupting the overall cellular composition of either tissue (**Fig. 2C**). In bone marrow, neutrophils were markedly expanded, accompanied by pronounced reductions in mature B cells and T cells. Other lymphoid populations, including developing and pre-B cells, were also decreased, whereas HSC abundance remained relatively stable (**Fig. 2C**). In the spleen, neutrophils and macrophages were increased, while both CD4 and CD8 T cell populations were reduced. B cell subtypes were affected differently, with follicular B cell population increasing in response to KO and while marginal zone B cell population expanded (**Fig. 2C**). Notably, HSC population size remained relatively stable in the KO, and the emergence of neutrophil precursor populations was observed, consistent with the extramedullary myelopoiesis and corroborating the immunophenotype previously reported(17). Together, these results indicate the CK2α loss does not abolish hematopoietic organization but induces a myeloid-skewed, lymphoid-depleted phenotype with prominent immune remodeling across tissues.

### HSCs exhibit the strongest transcriptional response to CK2α loss

Following cell clustering and cell type annotations, to identify transcriptional differences within cell types between WT and KO, we performed differential gene expression analysis (**Fig. 3A and B, Tables S1 and S2**) using edgeR (33). For both tissues, HSCs had the largest transcriptional response in KO. Among the most significantly differentially expressed genes were those involved in innate immune signaling, inflammation, and metabolic stress (**Fig. 3C and D**).

**Figure 3.**
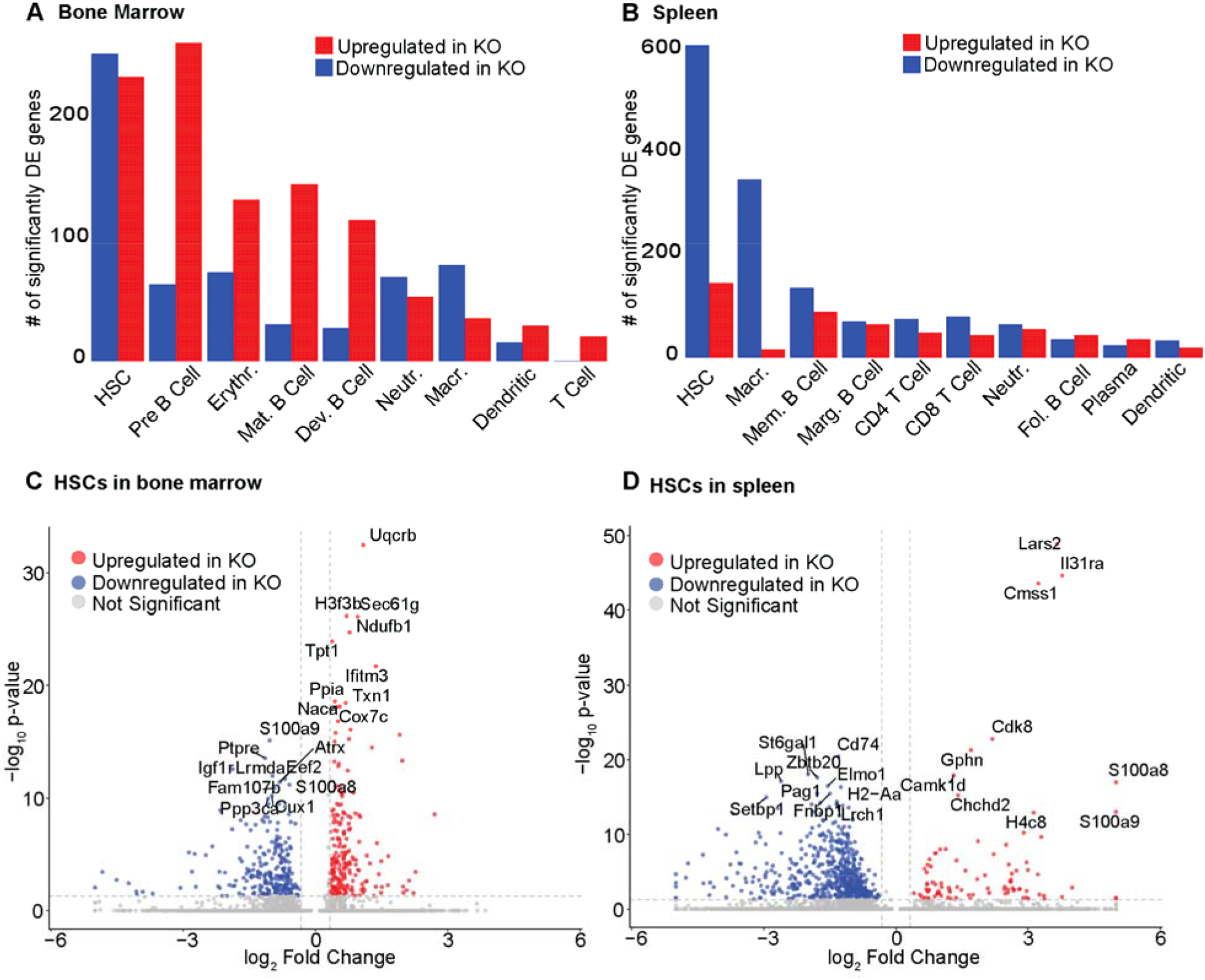
Differential gene expression analysis. (**A and B**) Number of significantly upregulated and downregulated genes per cell type in (**A**) bone marrow and (**B**) spleen (25% fold change, adjusted p-value less than 0.05). (**C and D**) Volcano plots showing significantly differentially expressed genes in the HSCs of (**C**) bone marrow and (**D**) spleen. The top 10 upregulated and downregulated genes ranked by smallest adjusted p-value are indicated.

Two of the top-ranked differentially expressed genes (DEGs) are shared between tissues, and notably, display opposing expression patterns between BM and SPL. *S100a8* and *S100a9* encode calcium-binding proteins that function as pro-inflammatory “alarmins”, a class of endogenous danger-associated molecular patterns (DAMPs) released or upregulated in response to cellular stress or tissue injury (34). Alarmins act as early warning signals that activate innate immune pathways by engaging pattern recognition receptors, thereby promoting cytokine production and inflammatory signaling (34). In bone marrow HSCs, the gene expression of these two alarmins was strongly downregulated in response to CK2α deletion (**Fig. 3C**), suggesting suppression of these stress-associated inflammatory mediators within the bone marrow niche. In contrast, these two genes were strongly upregulated in spleen HSCs (**Fig. 3D**), indicating that CK2α loss elicits tissue-specific immune activation programs and highlighting divergent inflammatory states between the two compartments.

Beyond the S100 family, the most significant DEGs further characterize a shift toward metabolic stress and immune priming in CK2α-deficient HSCs. In the bone marrow, upregulated genes were heavily enriched for components of the mitochondrial respiratory chain, including *Uqcrb, Ndufb1*, and *Cox7c* (**Fig. 3C**), consistent with increased reliance on oxidative phosphorylation and potential mitochondrial stress (35). The antioxidant gene *Txn1* was also significantly upregulated (**Fig. 3C**), suggesting activation of redox-buffering mechanisms in response to elevated reactive oxygen species (ROS) (36). Additionally, the interferon-induced gene *Ifitm3*, which restricts viral entry and reflects heightened innate immune signaling (37), was among the most upregulated genes, while the growth and survival receptor *Igf1r* was downregulated, pointing to a loss of normal homeostatic signaling.

In the spleen HSCs, transcriptional changes were characterized by heightened immune receptor expression coupled with impaired antigen presentation capacity. The cytokine receptor *Il31ra*, which mediates inflammatory and immune-modulatory signaling (38), was significantly upregulated (**Fig. 3D**), further supporting an immune-activated phenotype in this tissue. Conversely, genes essential for major histocompatibility complex class II (MHC II)-mediated antigen presentation (39), including *Cd74* and *H2-Aa*, were markedly downregulated (**Fig. 3D**), suggesting compromised interactions with adaptive immune cells. Collectively, these gene-level changes indicate that CK2α loss drives HSCs away from a quiescent, homeostatic state and toward a stressed, immune-primed phenotype in a tissue-specific manner.

### CK2α-deficient HSCs are activated for immune, metabolic, and stress-response pathways

To determine the functional consequences of CK2α-dependent transcriptional changes, we performed pathway analysis for the HSC populations in bone marrow and spleen samples using AUCell (40) and identified the differentially activated hallmark pathways in KO samples when compared to WT control (**Fig. 4A and B, Table S3**). Across both tissues, CK2α-deficient HSCs were activated for metabolic and stress-associated pathways, whereas they were deactivated for signaling pathways associated with stem cell maintenance and regulatory balance. At the same time, we also observed distinct activation and deactivation patterns in the two tissues in response to KO, indicating that CK2α loss reshapes HSC regulatory programs in a niche-dependent manner (**Fig. 4A and B**).

**Figure 4.**
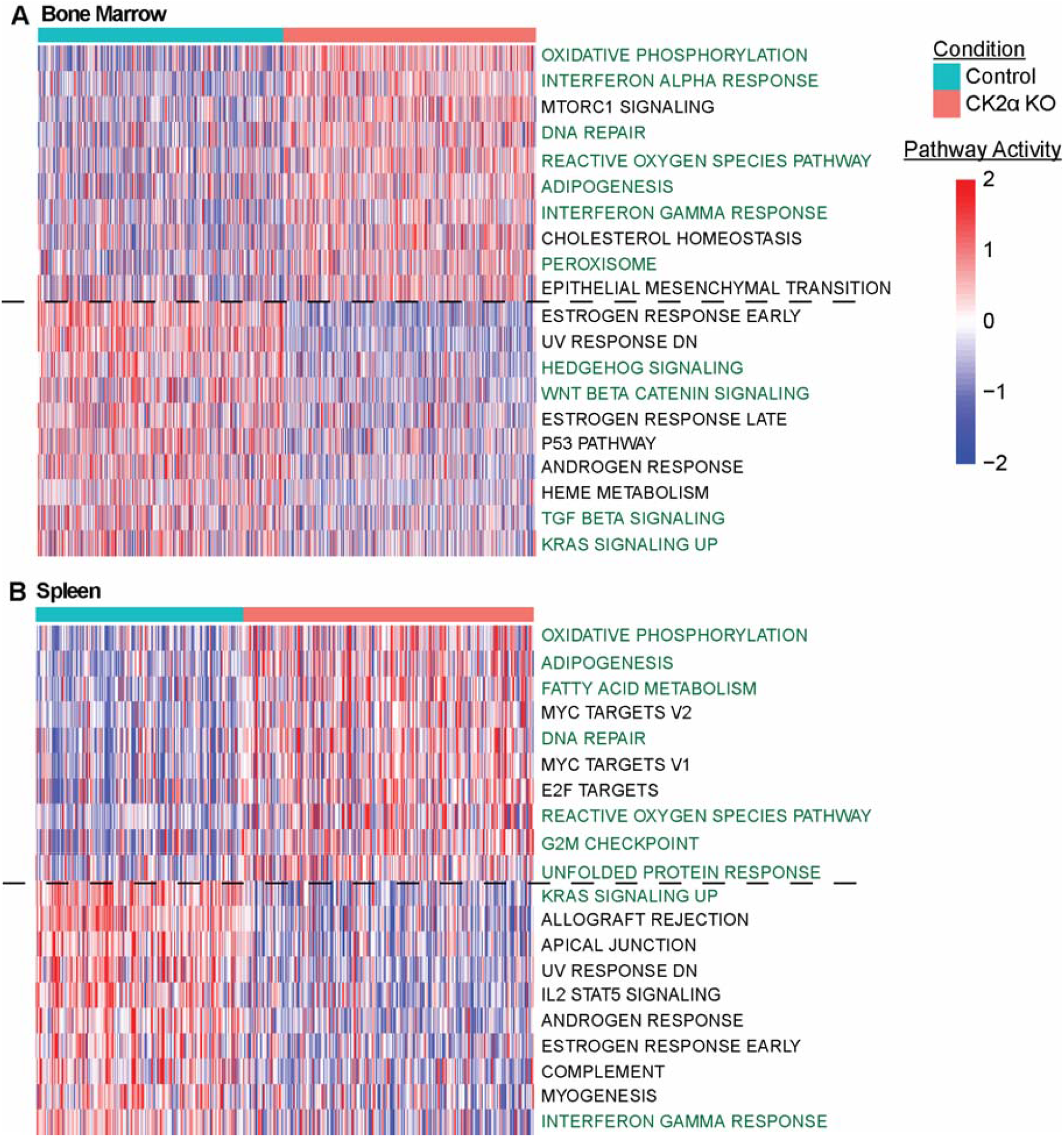
AUCell pathway analysis of differentially expressed genes in HSCs. Top 10 upregulated and downregulated hallmark gene sets by adjusted p-value in (**A**) bone marrow and (**B**) spleen.

In bone marrow, when compared to WT control, CK2α-deficient HSCs were activated for oxidative phosphorylation, adipogenesis, and peroxisome pathways (**Fig. 4A**), indicating increased mitochondrial and lipid-associated metabolic activity. This outcome aligns with the differential upregulation of mitochondrial respiratory chain genes such as *Uqcrb, Ndufb1*, and *Cox7c*, along with the antioxidant gene *Txn1* (**Fig. 3C**), suggesting elevated oxidative metabolism with compensatory redox regulation. Such metabolic activation contrasts with the low-metabolic state typically associated with quiescent stem cells (41). Bone marrow CK2α-deficient HSCs were also activated for reactive oxygen species and DNA repair pathways (**Fig. 4A**), consistent with increased oxidative and genotoxic stress.

Immune signaling pathways were also activated in CK2α-deficient bone marrow HSCs compared to control. Both interferon alpha and interferon gamma response pathways were elevated in response to CK2α-deficiency (**Fig. 4A**), corresponding with increased expression of the interferon-stimulated gene *Ifitm3* (**Fig. 3C**). These signatures indicate activation of interferon-associated transcriptional programs following CK2α loss. Together, these immune and stress-related pathways suggest that bone marrow HSCs adopt a more immune-responsive transcriptional state when CK2α signaling is disrupted.

When compared to WT control, CK2α-deficient bone marrow HSCs were downregulated for developmental signaling pathways associated with stem cell maintenance and lineage regulation (**Fig. 4A**). Specifically the Wnt/β-catenin signaling, Hedgehog signaling, TGFβ signaling, and KRAS signaling pathways are widely associated with HSC self-renewal and niche-dependent signaling (42), suggesting that CK2α deficiency may possibly affect transcriptional programs that maintain homeostatic stem cell functions.

In the spleen sample, CK2α-deficient HSCs were similarly activated for oxidative phosphorylation, adipogenesis, and fatty acid metabolism pathways (**Fig. 4B**), indicating a shift toward increased metabolic activity. Stress-associated pathways were also activated, including reactive oxygen species, DNA repair, unfolded protein response, and G2/M checkpoint pathways (**Fig. 4B**), indicating increased oxidative, proteotoxic, and proliferative stress in these cells. These signatures suggest increased genotoxic, oxidative, and proteotoxic stress, consistent with the observed metabolic reprogramming. Such pathways are commonly activated in HSCs under conditions of chronic stress or inflammation (43). Consistent with this immune-associated transcriptional environment, splenic CK2α-deficient HSCs showed increased expression of the alarmins *S100a8* and *S100a9* (**Fig. 3D**), genes commonly associated with innate immune activation.

The splenic CK2α-deficient HSCs were upregulated for KRAS signaling and interferon gamma response pathways when compared to wild type control (**Fig. 4B**). These pathways indicate engagement of baseline signaling and immune-associated regulatory programs in the absence of CK2α disruption. CK2α-deficient HSCs also showed reduced expression of antigen presentation genes such as *Cd74* and *H2-Aa* (**Fig. 3D**), suggesting altered immune interaction capacity in these cells.

Together, these results demonstrate that CK2α loss shifts HSC transcriptional programs toward metabolic activation, immune signaling, and cellular stress responses. At the same time, it attenuates the signaling pathways associated with developmental regulation and stem cell homeostasis. These pathway modulations support a model in which CK2α is associated with maintaining homeostasis within HSCs while limiting stress-associated and immune-driven transcriptional reprogramming.

### Immune-associated regulators emerge as hubs in CK2α-deficient HSCs

To identify regulatory rewiring events underlying the transcriptional reprogramming observed in response to CK2α KO, we employed gene regulatory networks (GRNs), which model transcription factor (TF)-to-target gene interactions. For this purpose, using CellOracle (27), we inferred GRNs for the HSC populations in the wild type and CK2α-deficient samples in the bone marrow and spleen, and compared the centrality degrees of the transcriptional regulators between the two genotypes (**Table S4**). In bone marrow HSCs lacking CK2α, immune- and stress-responsive TFs, including *Nfkb1, Hes1*, and AP-1 family members (*Fos* and *Jun*) emerged as network hubs, whereas TFs associated with lineage stabilization, such as *Junb*, showed reduced regulatory influence based on both rank (**Fig 5A and S5A**). Being a key mediator of inflammatory and survival signaling (44), the increased centrality of *Nfkb1* in the CKα-deficient HSCs is consistent with the activation of interferon subset and immune response pathways observed in CK2α-deficient HSCs. Similarly, increased influence of *Fos* and *Jun* suggests enhanced reliance on AP-1-driven transcriptional programs activated by stress, cytokine, and reactive oxygen species signaling.

**Figure 5.**
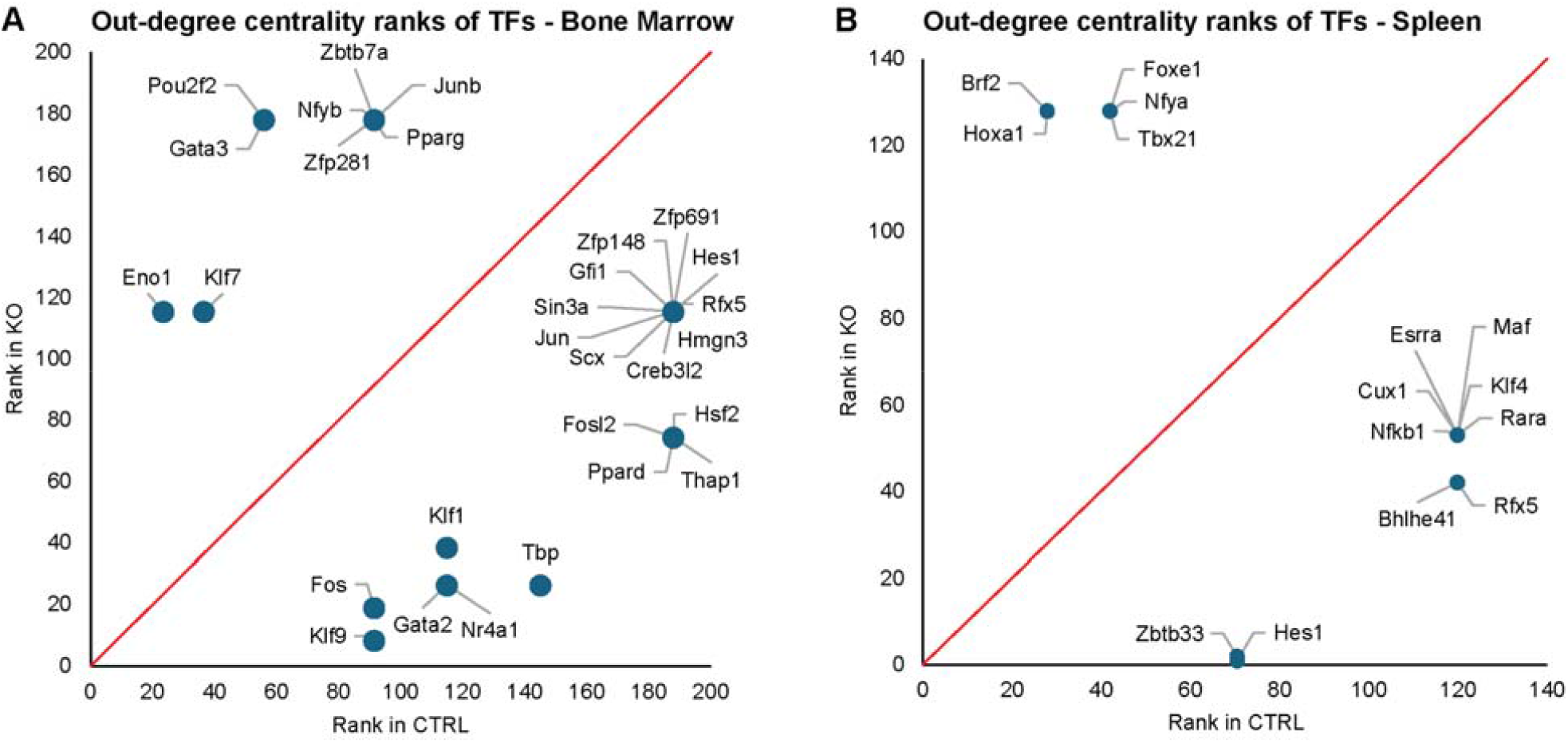
Out-degree centrality ranks for the regulator TFs in the respective GRNs in bone marrow (**A**) and spleen (**B**). Ranks for the same TFs in control and knockout are shown in both plots. Lower numbers in the ranks indicate higher centrality. The lower triangle of each plot shows the TFs that have larger centrality degrees in KO whereas the top triangle shows the TFs with reduced influence in KO.

The emergence of *Hes1* as a key regulator further implicates altered Notch-associated transcriptional control in response to CK2α loss. *Hes1* has been shown to restrain differentiation and reinforce progenitor-associated transcriptional states (45), and its increased network influence may contribute to the maintenance of an immature regulatory profile despite widespread transcriptional changes. In contrast, *Junb*, an AP-1 family member known to stabilize regulatory circuits involving TGF-β and Notch signaling and to limit excessive progenitor activation (46), exhibited reduced centrality, indicating a qualitative shift in AP-1 usage away from homeostatic control toward stress-responsive transcriptional regulation. This redistribution of AP-1 activity parallels the observed loss of growth factor and homeostatic pathway activation in CK2α-deficient HSCs.

In the spleen HSC population, a related but distinct pattern of regulatory rewiring was observed. *Nfkb1* and the immune-associated regulator *Rfx5* showed increased centrality following CK2α loss (**Fig 5B and S5B**). *Rfx5* is required for MHC class II gene expression (47), linking network changes to the downregulation of antigen presentation genes such as *Cd74* and *H2-Aa* observed at the transcription level. Together with the upregulation of inflammatory mediators such as *S100a8* and *S100a9* in splenic HSCs (**Fig. 3D**), these findings suggest that CK2α loss promotes consolidation of regulatory control around immune-associated transcriptional circuits in this tissue.

Collectively, network rewiring events indicate that CK2α loss shifts regulatory control away from lineage-associated transcription factors and toward immune- and stress-responsive transcription factors. This shift parallels the activation of interferon, oxidative stress, and DNA damage response pathways identified in pathway analysis and implies a mechanistic link between transcription factor activity and observed gene expression changes. Thus, CK2α appears to contribute to maintaining balanced regulatory input across immune, stress, and lineage-associated programs, and its absence biases regulatory control toward transcription factors that coordinate inflammatory and stress-associated transcriptional responses.

### Intercellular immune signaling is rewired upon CK2α loss

To understand the changes triggered by CK2α loss in the intercellular ligand-receptor between the HSCs and the immune cell types, we performed crosstalk analysis using LIANA+ (48). This analysis suggested that CK2α loss reshapes immune- and stress-associated signaling between HSCs and their microenvironment. In the bone marrow, CK2α-deficient mice showed increased signaling from HSCs to macrophages through the ligand-receptor pairs *Fn1-Itga4* and *Fn1-Itgb1* (**Fig. 6A**). Since *Itga4* and *Itgb1* together form the VLA-4 integrin complex, these interactions indicate enhanced FN1-VLA-4 signaling from HSCs to macrophages in the deficiency of CK2α. FN1 engagement of VLA-4 induces the ERK1/ERK2 pathway (49,50), which promotes macrophage activation, inflammatory cytokine production, and stress response programs. Thus, the emergence of this signaling axis is consistent with the increased inflammatory and stress-responsive transcriptional programs observed in CK2α-deficient bone marrow HSCs and suggests that loss of CK2α promotes a feed-forward interaction between HSCs and macrophages that may reinforce inflammatory niche remodeling.

**Figure 6.**
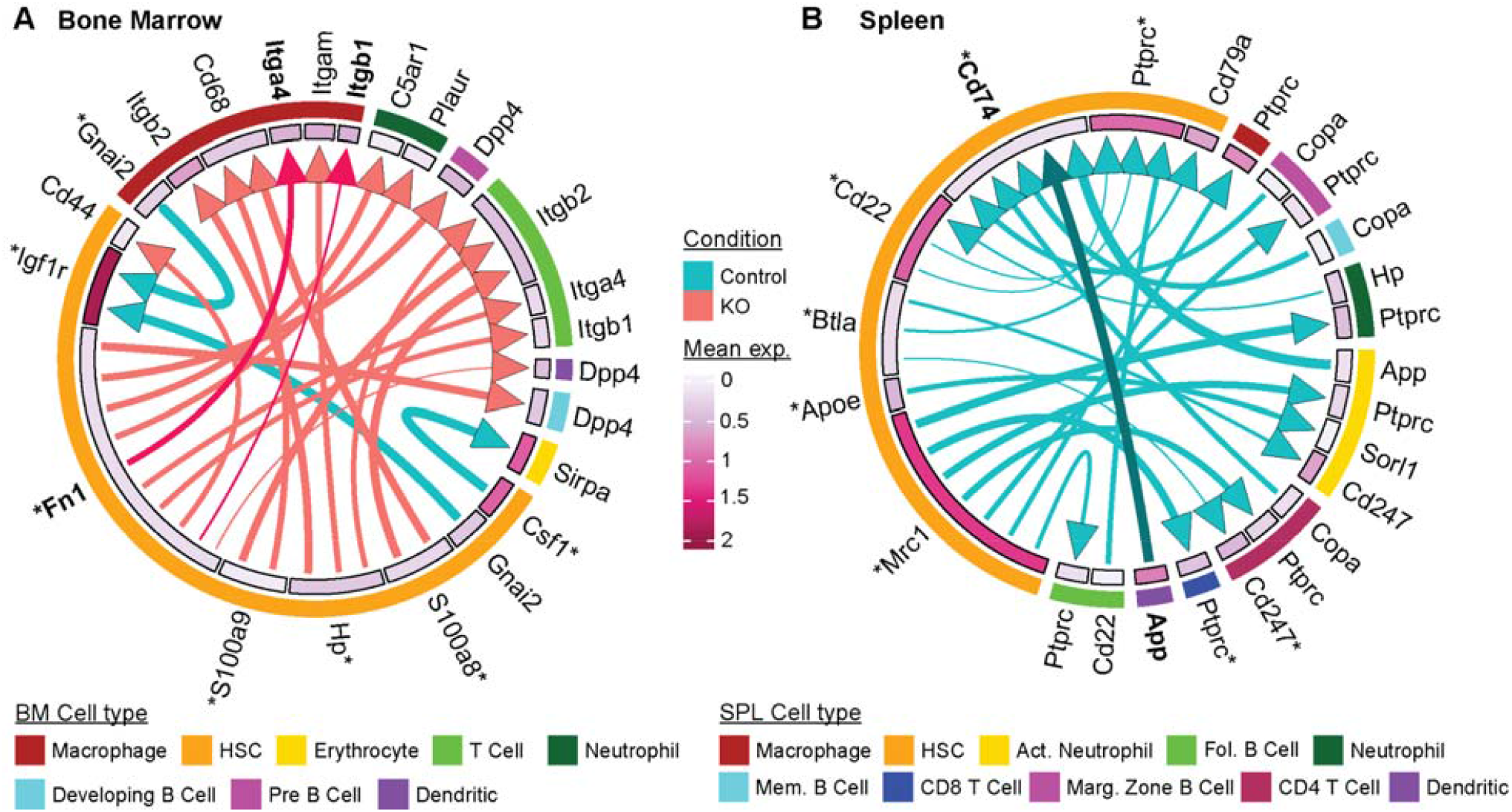
Intercellular crosstalk between HSCs and immune cell types. Circos plots indicating the ligand and receptor interactions for which either the ligand or receptor is differentially expressed in bone marrow (**A**) and spleen (**B**). Top 20 interactions are shown for each tissue, based on the magnitude rank. Arrow thickness indicates the significance of the pair. An asterisk indicates a significantly differentially expressed gene in KO.

In contrast, splenic cell communication analysis identified an *App-Cd74* interaction from dendritic cells to HSCs that was present in control mice but lost following CK2α deletion (**Fig. 6B**). CD74-mediated signaling has been linked to activation of ERK1/ERK2 and MAPK pathways (51), as well as regulation of antigen presentation and cellular stress responses (52). Loss of APP-CD74 signaling in the knockout therefore suggests impaired dendritic cell-mediated communication with HSCs, potentially contributing to the reduced expression of antigen presentation genes and altered immune regulatory state observed in CK2α-deficient splenic HSCs. Together, these findings indicate that CK2α loss not only rewires intrinsic transcriptional programs within HSCs, but also alters the extracellular signaling networks that connect HSCs with surrounding immune cells.

## Discussion

Our study reveals a previously unappreciated role for CK2α in coordinating immune and transcriptional programs within HSCs in a tissue-dependent manner. Using single-cell RNA sequencing of bone marrow and spleen from hematopoietic-specific CK2α knockout mice, we demonstrate that while overall lineage composition is largely maintained, HSCs exhibit the most pronounced transcriptional perturbations following CK2α loss. This heightened sensitivity identifies the HSC compartment as a primary nexus of molecular dysregulation, where the absence of CK2α triggers a shift away from homeostatic maintenance toward a state of chronic stress.

A key finding of our work is the immune-centered transcriptional rewiring of HSCs in response to CK2α deletion. By combining our differential expression and pathway analysis results, we observe a coordinated transition toward an immune-primed state across both tissues. Differential expression analyses revealed strong dysregulation of immune and inflammatory genes such as *S100a8* and *S100a9*, and interferon-responsive transcripts. Interestingly, several of these genes, including *S100a8* and *S100a9*, exhibited opposite regulation in bone marrow versus spleen, suggesting that CK2α-dependent immune control is highly context specific. This divergent response underscores the fact that the local microenvironment, such as the differing oxygen levels or vascular structures of the bone marrow and spleen, actively shapes how stem cells interpret the loss of kinase signaling.

Pathway and network analyses further clarify the organization of these transcriptional changes. CK2α-deficient HSCs are activated for interferon signaling, reactive oxygen species (ROS) pathways, and metabolic pathways, alongside emergent regulatory hubs including *Nfkb1, Hes1, Rfx5*, and AP-1 family members. The increase in the centrality for these regulators in the KO sample HSC population indicates that loss of CK2α concentrates regulatory influence onto a smaller set of stress- and immune-associated transcription factors. The transition from a distributed network of lineage-stabilizing factors, such as *Junb*, toward a concentrated influence in stress-responsive hubs like *Nfkb1* provides a structural explanation for the loss of lineage fidelity (46). This concentration of influence may prioritize stress-responsive outputs over homeostatic balance, potentially limiting the ability of HSCs to buffer against additional perturbations like chronic inflammation or aging.

Joining the results from the differential expression of genes, pathway analysis, GRN inference, and cell-cell communication supports a model in which CK2α loss promotes distinct niche-driven stress programs in bone marrow and spleen (**Fig. 7**). In the bone marrow, CK2α-deficient HSCs gain FN1-VLA-4 signaling to macrophages through the *Fn1-Itga4/Itgb1* axis, potentially activating ERK-dependent cytokine production and reinforcing inflammatory signaling within the niche (53). This model is supported by the emergence of Fos as a central regulator TF in the knockout GRN, where it is predicted to directly activate the expression of the *Fn1* ligand in HSCs. Fn1 then can bind to the VLA-4 complex on macrophages and promote the downstream signaling cascade (54), leading to the immune-primed, stress-responsive phenotype inferred from the pathway analysis. In the spleen, CK2α-deficiency leads to the loss of the *App-Cd74* axis between dendritic cells and HSCs. This is supported by the emergence of *Rara* as a regulator TF in the knockout GRN, with a negative coefficient regulating *Cd74* expression in HSCs. The CK2α-deficient HSCs therefore lose the expression of *Cd74*, leading to a loss of the *App-Cd74* signaling axis. *Cd74* is essential for MHCII antigen presentation (39) and likely contributes to the downregulation we see of antigen presentation genes in the splenic HSCs. *App-Cd74* signaling has also been linked to the promotion of a more quiescent cell state (55), and its loss would be expected to exacerbate the metabolically-disrupted stress-responsive pathways which are activated in the spleen.

**Figure 7.**
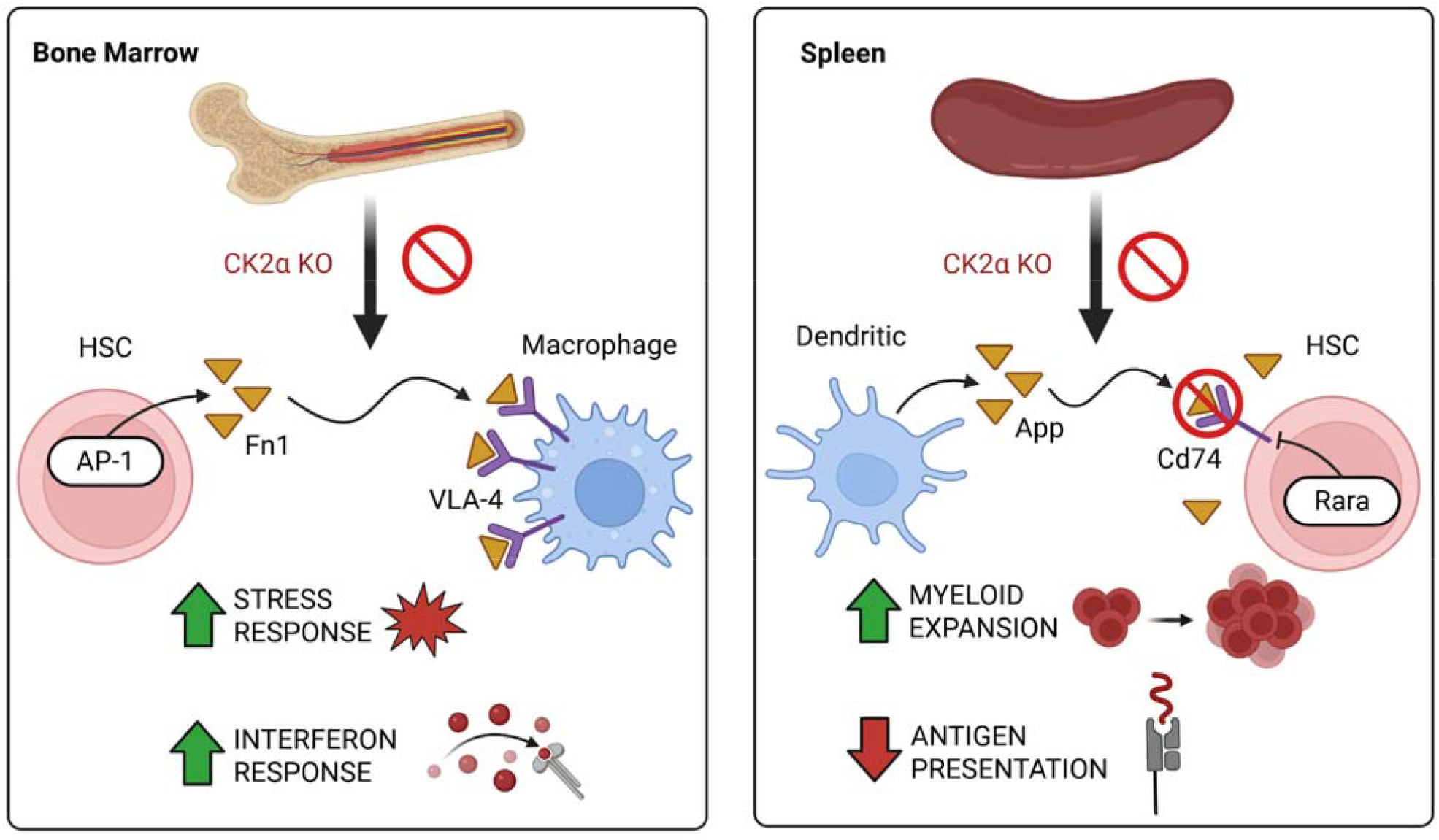
Summary of the main findings in this study. Differential activation of notable intercellular communicated and intracellular pathways as a response to CK2α KO, with the central focus on the HSC population are illustrated for both tissues.

In our single-cell data, we successfully mapped 11 distinct cell types in the bone marrow and 12 in the spleen, including various B cell, T cell, and myeloid subsets. Future studies leveraging this resource can explore whether the systemic myeloid expansion and lymphoid depletion are driven solely by stem-cell-level priming or if downstream progenitors and mature immune cells undergo their own distinct forms of tissue-specific transcriptional rewiring. Expanding our analytical framework to these other populations will be essential to fully resolve the systemic impact of CK2α loss.

These findings also have important potential clinical significance. CK2α inhibitors are under investigation in cancer therapy, including hematologic malignancies, where synthetic kinase inhibition using clinical grade small modlecule inhibitor, CX-4945, selectively targets leukemic cells sparing normal blood cells (15,56–58). Our results suggest that CK2α inhibition could exert tissue-specific effects on healthy HSCs, potentially altering immune and inflammatory programs in a context-dependent manner. Consequently, therapeutic strategies targeting CK2 may need to account for these specific effects on the steady-state immune landscape of the bone marrow and spleen.

These findings have important therapeutic implications for Immunotherapy. The systemic rewiring following CK2 inhibition presents a potent synergistic opportunity when combined with immunotherapy. By pharmacologically targeting CK2α (ex: CX-4945), we may strategically leverage this immune-priming effect to enhance the production and mobilization of immune-responsive progeny. Such an approach could augment the efficacy of checkpoint inhibitors or adoptive cell therapies by lowering the activation threshold of the hematopoietic compartment and intensifying the innate inflammatory response against malignant cells. Future studies must determine if the “immune-primed” stress state observed in healthy HSCs translates to leukemic stem cells.

In conclusion, our study identifies CK2α as a key regulator of immune and stress-response transcriptional programs in HSCs, with distinct effects in bone marrow versus spleen. These results expand our understanding of CK2’s role beyond general cell survival and proliferation to include niche-dependent modulation of immune gene expression and transcriptional network architecture. By linking kinase activity to tissue-specific immune regulation in stem cells, our findings provide a framework for exploring both normal hematopoiesis and disease-associated perturbations of HSC function.

## Methods

### Animals

To investigate the involvement of CK2α in shaping hematopoietic transcriptional states, we used a hematopoietic-specific CK2α conditional knockout (KO) mouse model generated by crossing Vav-iCre mice with *Csnk2a1* floxed mice (**Fig. 1A)**. It produced Vav iCre Csnk2a1^fl/fl^ (KO) and Vav iCre Csnk2a1^+/+^ (WT) offsprings. CK2α deletion was confirmed with transnetyx genotyping at the genetic level (**Fig. S1**) and with Western blot at the protein level (**Fig. S2**) in bone marrow (BM) and spleen (SPL). Mice with floxed *Csnk2a1* alleles were kindly gifted from Drs. Flajolet and Rebholz (City College of New York, New York, NY) (59). Vav-iCre mice that express improved Cre recombinase (iCre) under the control of mouse vav 1 (Vav1) promoter were purchased from The Jackson Laboratory (60). These mice were crossed on a C57BL/6 background for at least three generations. The wild type (WT) Vav-CreCK2α+/+, and knockout (KO) Vav-iCreCK2αf/f mice used in this study were both 8-week-old females. Littermates of the Vav-iCreCK2αf/f (KO) were used as wild-type controls in each specific experiment. The deletion of *Csnk2a1* was confirmed by genotyping performed using real time PCR (TransnetYX, Cordova, TN). Primers are listed in **Table S5**. All of the mice were bred and maintained in pathogen free conditions at Pennsylvania State University College of Medicine, Comparative Medicine facility. The protocols were approved by the Institutional Animal Care and Use Committee of the Pennsylvania State University.

The Vav-iCre conditional knockout strategy targets specific exons of the *Csnk2a1* (CK2α) locus to disrupt protein expression selectively in hematopoietic cells. Because exons outside the floxed region remain intact, transcripts corresponding to these exons can still be detected in RNA sequencing data from knockout cells. Therefore, residual CK2α RNA signal may be observed in knockout samples, although protein expression is effectively ablated in these cells as per the western blot (Fig. S#) of the CK2 protein subunits.

### Western Blotting

Whole cell lysate was prepared from BM cells Vav-iCreCK2α+/+, Vav-iCreCK2αf/f without RBC lysis using cell lysis buffer (Cell Signaling Technology). Initially, 5 µl of cell lysates from each sample were subjected to Western blot analysis with β-actin antibody (Cell Signaling), and band intensities were normalized. Proteins from normalized cell lysates were separated by 4-20% SDS-PAGE in Laemmli buffer and electro-transferred to a PVDF membrane. Further, the membranes were blocked with TBST (20 mmol/L Tris, pH 7.5 with 150 mmol/L NaCl and 0.05% Tween-20) containing 5% non-fat milk powder for 1h. Then, the blot was probed with primary antibodies specific for mouse CK2α (E-7 clone, Santa Cruz; SC-373894) overnight at 4°C. Membranes were washed extensively (4x) using TBST and incubated with HRP conjugated secondary antibodies (Cell Signaling Technology) for one hour at room temperature. The blots were developed with an enhanced chemiluminescence substrate and visualized using a Bioimaging system (Bio-Rad). Mouse anti-CK2α antibody was used at 1:10000 dilution and secondary antibody was used at dilution of 1:3000.

### Cell collection for single cell RNAseq

The bone marrow and spleen cells from the Vav-iCreCK2α+/+ and Vav-iCreCK2αf/f mice were isolated on the same day by placing the spleen into a cell strainer (Fisherbrand). After crushing, the spleen cells were collected in the FACS buffer and washed 2 times by spinning at 1200 rpm for 5 minutes. The supernatants were then removed, and the cells were resuspended in 1 mL of RBC lysis buffer (eBioscience) for 8 minutes. Following resuspension, FACS buffer was added and spun at 1200 rpm again for 5 mins. After spinning, the supernatants were removed, and the cells were resuspended in 1mL of FACS buffer. For bone marrow, the bone was flushed with the FACS buffer to collect the cells. After that, the cells were washed with FACS buffer by spinning at 1200 rpm for 5 minutes. The supernatants were then removed, and the cells were resuspended in 1 mL of RBC lysis buffer (eBioscience) for 4 minutes. After that, FACS buffer was added and spun at 1200 rpm again for 5 minutes. After spinning, supernatants were removed, and the cells were resuspended in 1mL of FACS buffer. Cells from both tissues were counted using trypan blue, and 1×10^5^ good-quality cells, having >80% viability, <20% debris, and <15% clumps, were submitted to the Genome Sciences core for single cell RNAseq.

### scRNA-Seq library preparation and sequencing

Cell 3□ Reagent Kits v3.1 (Dual Index), following manufacturer’s protocol/ Briefly, 20000 cells were loaded into each well to capture 10000 cells. cDNA from each cell was barcoded and sequencing libraries prepared using the indices from 10x Genomics Dual Index Plate TT Set A (PN-3000431). 11 cycles for cDNA amplification and 13 cycles for sample index PCR were carried out. Final libraries were assessed for size distribution and concentration using BioAnalyzer High Sensitivity DNA Kit (Agilent Technologies). Libraries were prepared in the Penn State College of Medicine Genome Sciences core (RRID:SCR_021123).

The libraries were pooled and sequenced on Illumina NovaSeq 6000 (Illumina), to get on average 200 million, paired end (Read 1 28 bp and Read 2 90 bp) reads, according to the manufacturer’s instructions. Samples were demultiplexed using bclconvert software (Illlumina). Detailed sequencing and alignment metrics are provided (**Table S6**). Adaptors were not trimmed during demultiplexing. Sequencing was done in the Penn State College of Medicine Genome Sciences core (RRID:SCR_021123).

### Quality control

Quality control (QC) was performed to remove low-quality cells, technical artifacts, and potential doublets prior to downstream analysis. Cells were filtered based on three primary metrics: (1) the total number of expressed genes, (2) the total number of UMI counts, and (3) the proportion of reads mapping to mitochondrial genes. Cells with fewer than 500 expressed genes or fewer than 1,000 UMI counts, indicative of empty droplets or poor library complexity, were excluded. Cells with more than 10,000 expressed genes or more than 20,000 UMI counts were removed as potential doublets or multiplets. To exclude damaged or dying cells, cells with >10% mitochondrial RNA content were filtered out. Genes detected in fewer than 3 cells were excluded from downstream analyses to reduce noise from low-abundance transcripts.

To identify and remove doublets more rigorously, we applied a doublet-detection algorithm (DoubletFinder), using an expected doublet rate of 7.5% and parameter settings of pN = 0.25, pK = 0.09, and an nExp value proportional to the expected number of doublets based on recovered cell counts. Detected double cells were removed from the dataset before normalization. Each sample was randomly downsampled to 7,025 cells, matching the smallest sample, to ensure equal cell representation across biological replicates and to minimize sample-size-driven bias in downstream integration and clustering. All QC steps were carried out on raw count matrices prior to merging.

### Data normalization, scaling, dimensionality reduction, and clustering

All data processing and analyses were performed using Seurat v5.1.0 (30) in R. Normalization was performed using NormalizeData, applying log-normalization to total UMI counts. Gene expression values were then scaled and centered using ScaleData. The top 2,000 highly variable genes were identified using FindVariableFeatures.

Principal component analysis (PCA) was performed for dimensionality reduction using RunPCA, and the top 10 principal components (PCs) were retained for downstream analyses. Dimensionality reduction was performed using UMAP via RunUMAP, also using these top 10 PCs.

A shared nearest-neighbor graph was constructed with FindNeighbors, and clustering was performed with FindClusters using the Louvian algorithm at a resolution of 0.05, which yielded biologically interpretable and non-fragmented clusters. Control and knockout samples were processed together for each tissue to ensure joint embedding and direct comparison of cell states.

### Cell type annotation

Cell type identities were assigned using a combination of canonical marker gene expression and cluster-specific differential expression. First, the expression levels of the well-established mouse hematopoietic marker genes were assessed across clusters to assign preliminary labels. For clusters in which canonical marker expression did not clearly distinguish a cell type, cluster-specific marker genes were identified using FindAllMarkers and compared with reference signatures from Haemopedia (31).

Candidate identities were assigned based on concordance between (1) canonical marker expression, (2)genes upregulated in the clusters, and (3) Haemopedia lineage profiles. Final annotations were reviewed manually to ensure consistency with known hematopoietic biology in mouse bone marrow and spleen.

### Differential expression and pathway analysis

Differential expression gene (DEG) analysis was performed within each annotated cell type to compare control versus CK2α knockout conditions. Only cell types containing at least 10 cells per condition were retained for testing. For all eligible cell types, DEG was performed using FindMarkers with Wilcoxon rank-sum testing. To limit computational load and ensure balanced comparisons, a maximum of 250 randomly selected cells per condition were included in each test. Significant DEGs were defined as those with |log_2_ fold change| less than or equal to 0.322 (corresponding to a 1.25-fold change) and adjusted p-value less than 0.05 (benjamini-Hochberg correction).

Pathway activation scores for individual HSCs were quantified using the AUCell (40) package in R. Gene expression matrices were extracted from the RNA assay using normalized expression values. Gene sets were obtained from the MSigDB mouse collection (v2025.1) (61,62), and only hallmark gene sets were retained for analysis. Gene sets were intersected with the expressed genes in each dataset, and empty sets were excluded. Per-cell gene rankings were generated using AUCell_buildRankings function with default parameters, and gene set activity was quantified using AUCell_calcAUC, which computes the area under the recovery curve for each gene set within each cell’s ranked gene list, yielding an AUC score representing relative pathway activity per cell (40).

### Gene regulatory network construction and comparisons

GRNs were constructed from single-cell RNA-sequencing data using the CellOracle (27) framework (v0.20.0). Processed Seurat objects for bone marrow and spleen containing annotated cell types and condition labels were converted to compatible format and imported into CellOracle as raw count matrices. A combined cluster-and-condition column was generated to define cell populations for GRN construction, ensuring that both genotype and cell-type identity were considered. Base GRNs were derived from publicly available mouse scATAC-seq datasets provided by CellOracle, and transcription factor information was imported to define candidate regulatory edges.

Dimensionality reduction was performed via principal component analysis, and K-nearest neighbor (KNN) imputation was applied to smooth gene expression across similar cells. GRNs were inferred for each cluster-condition unit using CellOracle’s linear modeling approach, generating weighted TF-target interactions. Raw networks were filtered to retain edges with absolute coefficient values above a defined threshold (2000) and significance (p < 0.01). Filtered networks were then used to calculate network scores, including out-degree centrality for each gene within each cell-type-specific network.

For downstream analyses, the GRNs were exported per cluster, and only edges with non-zero coefficients were retained. Multiple-testing correction was applied using the Benjamini-Hochberg method, and edges with adjusted p-values < 0.01 were considered statistically significant. Out-degree centrality metrics were computed for all TFs in control and knockout networks, and Δ-rank values were calculated to quantify rewiring of regulatory influence following CK2α deletion. TFs in the top 5% of absolute d-rank were designated as significantly rewired. These analyses provided a quantitative, cell-type-resolved view of CK2α-dependent regulatory network alterations in hematopoietic tissues.

### Cell communication (crosstalk) inference

Cell-cell ligand-receptor interactions were inferred using LIANA+ (48) in Python v3.13. The mouse consensus resource was used to identify potential interactions among the cell types present in bone marrow and spleen samples. Two complementary methods were employed: the rank aggregate method, which scores across multiple interaction scoring methods to produce a consensus ranking, and CellPhoneDB (63), used specifically to extract ligand and receptor expression proportion values. Only ligand receptor pairs expressed in at least 10% of cells were considered. Interactions from knockout (KO) and control conditions were merged using an outer join to retain all unique and shared interactions across conditions. A magnitude interaction threshold of <0.01 was used to define significant interactions, while scores >0.1 were considered non-significant. Then, the inferred interactions were compared to find the interactions that were gained, lost, or transitioned in specificity between KO and control conditions. Complete interaction results for HSC crosstalk are provided in **Tables S7-S10**.

## Acknowledgements

We would like to thank the Penn State College of Medicine Genome Sciences Core (RRID:SCR_021123). This work is supported by National Center for Advancing Translational Sciences (KL2 TR002015); Hyundai Hope on Wheels Scholar Grant; Four Diamonds Fund of the Pennsylvania State University College of Medicine; John Wawrynovic Leukemia Research Scholar Endowment (C.G.); St. Baldrick’s Foundation; Team Connor and Rally Foundation.

