## Supplemental Information for "Lineage-specific CK2α deletion reshapes the transcriptome of hematopoietic stem cells toward an immune-primed state"

### CK2$\alpha$ inhibition reshapes the transcriptional landscape of hematopoietic stem cells toward an immune-primed state

Hannah Valensi[1], Rajesh Rajaiah[1], Marudhu Shanmugam[1], Daniyal Muhammad[1], Upendar Golla[1], Katherine Mercer[1], Anush Karampuri[2], Sinisa Dovat[1,3], Chandrika Behura[1]*, Yasin Uzun[1,3]*

[1] Department of Pediatrics, Pennsylvania State University College of Medicine, Hershey, PA

[2] Huck Institutes of Life Sciences, Pennsylvania State University, University Park, PA

[3] Department of Molecular and Precision Medicine, Pennsylvania State University College of Medicine, Hershey, PA

* Correspondence

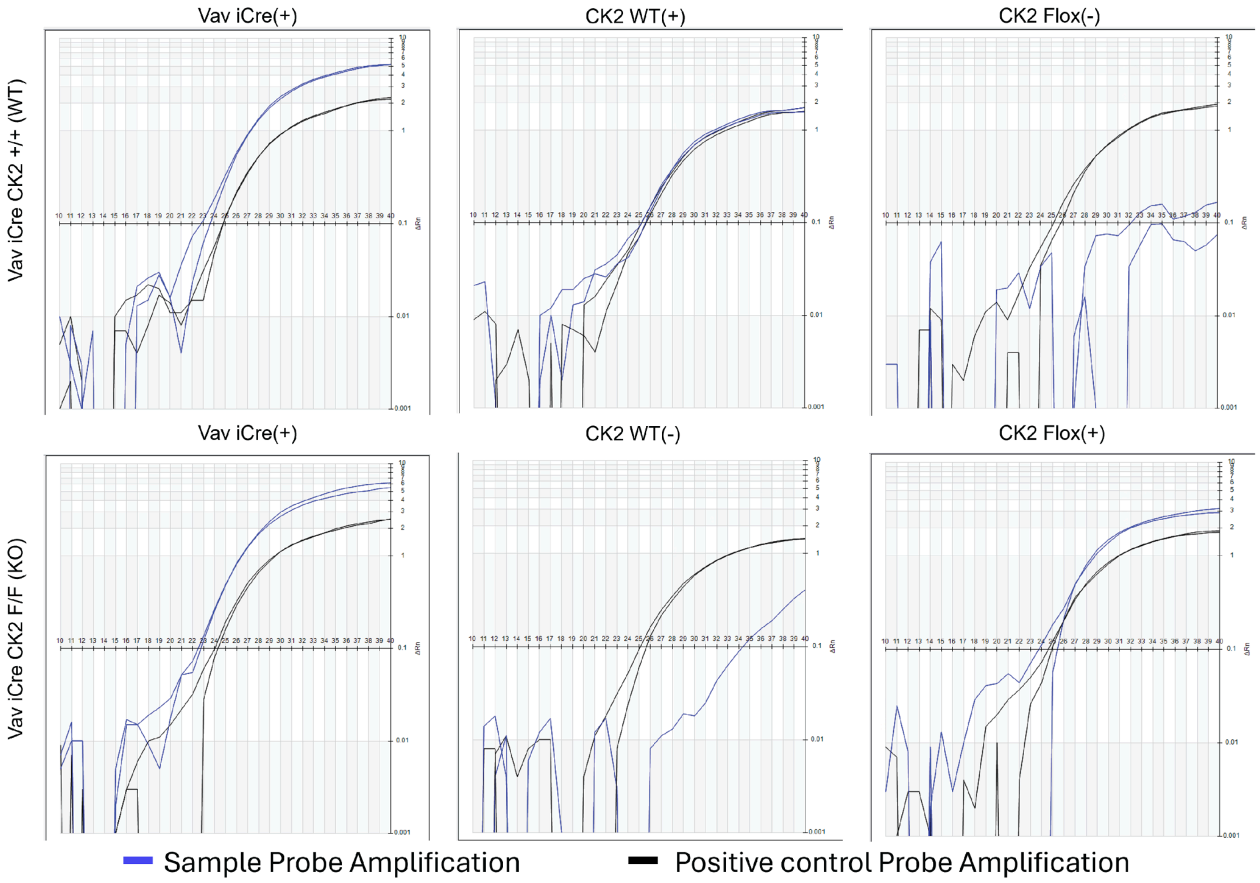


**Figure S1. Genotyping of Vav iCre CK2 conditional mice.** Genotyping results of tail end tissue from Transnetyx confirm Vav iCre CK2+/+ (WT) and Vav iCre CK2F/F (KO) mice. Blue curves represent amplification of the test mice for Vav iCre and CK2  WT and/or floxed allele, whereas black curves represent amplification of the positive control probe. Each curve shows fluorescence increase (ΔRn) across PCR cycles, with earlier cycle thresholds corresponding to positive allele detection. Automated qPCR assay depicts the Vav iCre transgene and distinguishes CK2 wild type and floxed alleles.

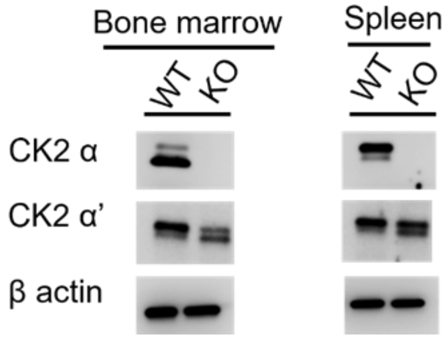


**Figure S2. Western blots of CK2 subunits for WT and CK2α KO mice.** Protein expression of CK2α and CK2α’ in WT (Vav-iCreCK2α+/+) and KO (Vav-iCreCK2f/f) mice bone marrow and spleen.

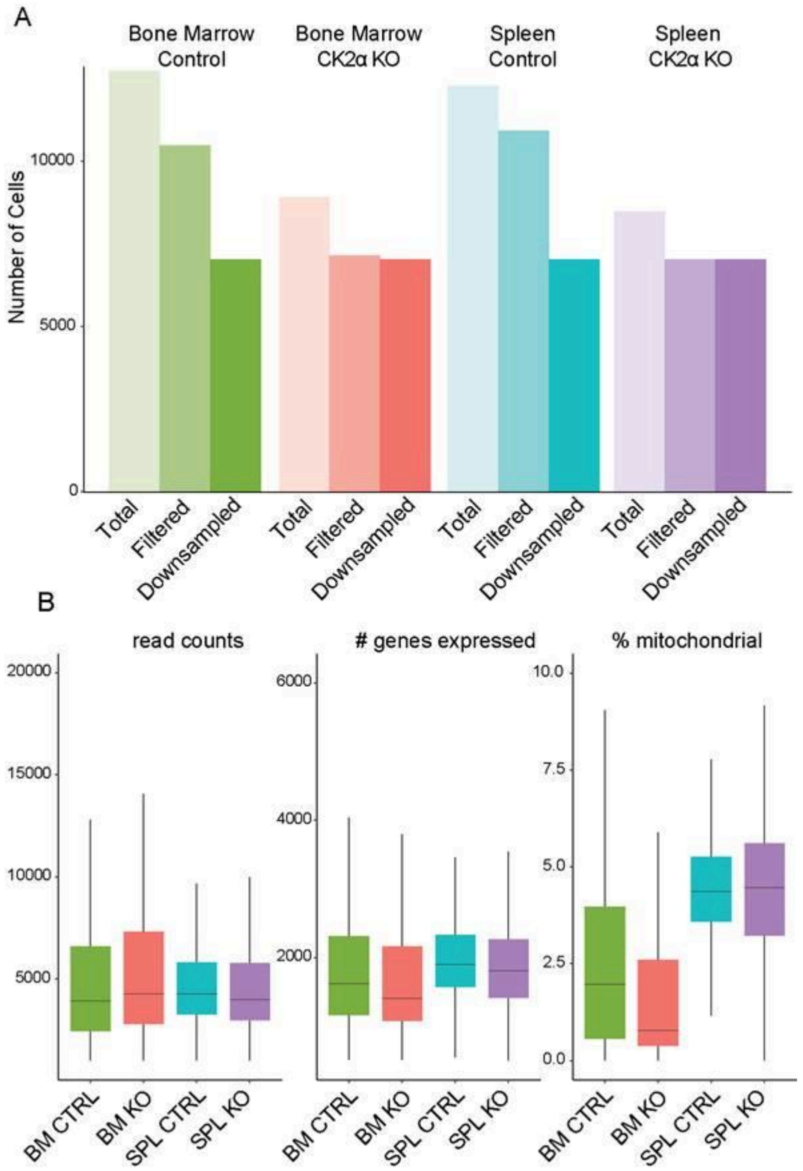


**Figure S3. Single cell quality control statistics.** (**A**) Bar plots showing cell counts per sample for total cells collected, cells surviving filtering criteria, and cells post-downsampling. (**B**) Box and whisker plots showing the number of read counts, genes expressed, and percent mitochondrial genes for each sample after filtering and downsampling.

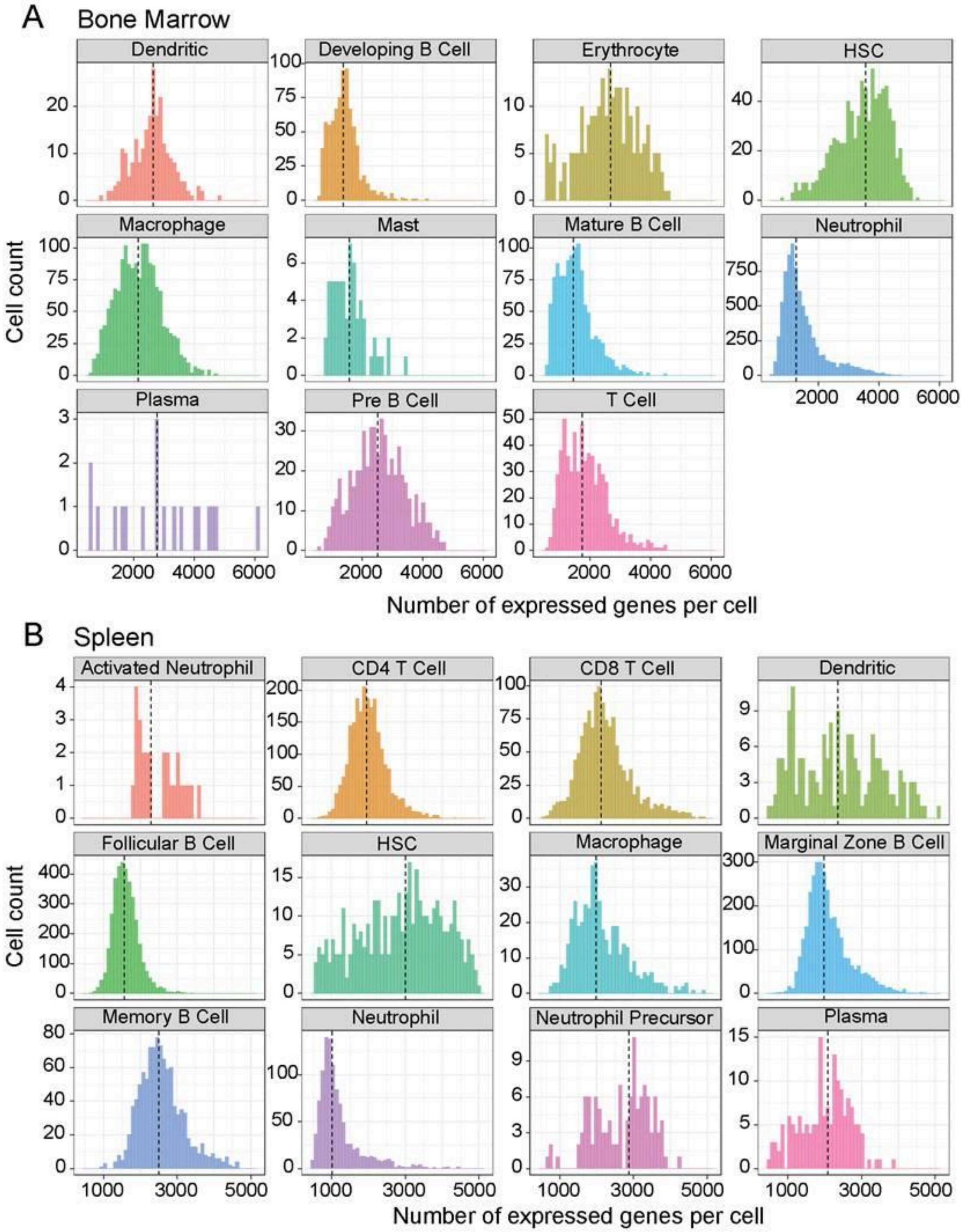


**Figure S4. Number of expressed genes per cell type.** Bar plots of the number of genes expressed per cell for each cell type in **(A)** bone marrow and **(B)** spleen tissues.

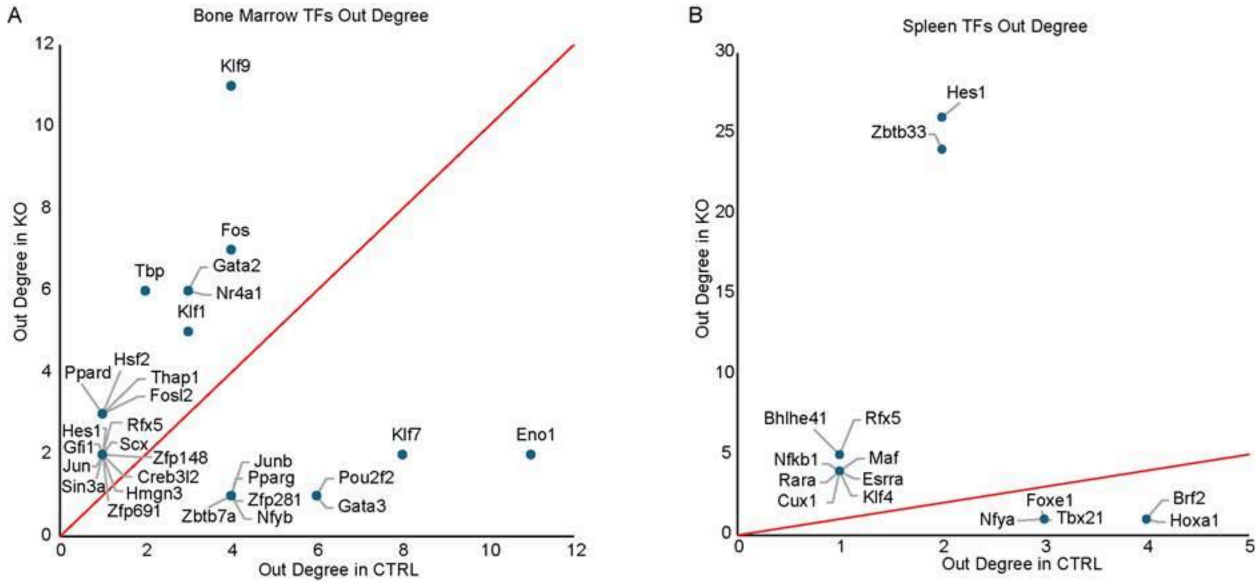


**Figure S5. Raw out-degree centrality scores.** Out-degree centrality scores for TFs in 95th percentile for delta out-degree. Out degree score for the same TFs in control and knockout in bone marrow (**A**) and spleen (**B**).

**Table S1: Differential expression from bone marrow:** This table contains differential gene expression results comparing KO and WT control cells in bone marrow. Separate sheets correspond to each annotated cell type. Columns include the gene symbol, unadjusted p-value, average $\log_2$ fold change, percentage of cells expressing the gene in control and KO samples, and adjusted p-value.

**Table S2: Differential expression from spleen:** This table contains differential gene expression results comparing KO and WT control cells in spleen. Separate sheets correspond to each annotated cell type. Columns include the gene symbol, unadjusted p-value, average $\log_2$ fold change, percentage of cells expressing the gene in control and KO samples, and adjusted p-value.

**Table S3: AUCell pathway:** This table contains AUCell pathway enrichment results comparing KO and WT control HSCs in bone marrow and spleen. Separate sheets correspond to the tissue. Only HALLMARK gene sets were analyzed. Columns include the gene set name, mean AUCell score in control and KO HSCs, fold change, $\log_2$ fold change, unadjusted p-value, and adjusted p-value.

**Table S4: Gene regulatory networks from HSCs:** This table contains inferred gene regulatory network interactions for HSCs in bone marrow and spleen under control and KO conditions. Separate sheets correspond to bone marrow control, bone marrow KO, spleen control, and spleen KO networks. Columns include the source transcription factor, target gene, mean interaction coefficient, absolute interaction coefficient, p-value, and negative $\log_{10}$-transformed p-value.

**Table S5: Primers used for the genotype of mice.**

| | Forward primer | Reverse primer | Probe |
|---|---|---|---|
| *Csnk2a1-FL* | ACATATTGACGCCTTTTTATACATGTCTCTT | TTCGTATAATGTATGCTATACGAAGTT | TTCAGAGGATATAATGGATAGGCT |
| *Csnk2a1-WT* | CAATGTCAAGAGTTACTTGGAATGTAGA | CTCTTTGACCACATCCTAACTATCCCTT | ACGCCTTTTTATACATGTCTCTT |
| iCre | TCCTGGGCATTGCCTACAAC | CTTCACTCTGATTCTGGCAATTTCG | ACCCTGCTGCGCATTG |

**Table 6. CellRanger Metrics**: This table includes the CellRanger metrics including; Estimated Number of Cells, Mean Reads per Cell, Median Genes per Cell, Number of Reads, Valid Barcodes, Sequencing Saturation, Q30 Bases in Barcode, Q30 Bases in RNA Read, Q30 Bases in UMI, Reads Mapped to Genome, Reads Mapped Confidently to Genome, Reads Mapped Confidently to Intergenic Regions, Reads Mapped Confidently to Intronic Regions,

Reads Mapped Confidently to Exonic Regions, Reads Mapped Confidently to Transcriptome, Reads Mapped Antisense to Gene, Fraction Reads in Cells, Total Genes Detected, Median UMI Counts per Cell for each sample.

**Table S7. Ligand-receptor interactions with bone marrow HSCs as source:** This table contains predicted ligand-receptor interactions in which bone marrow HSCs act as the source cell type. Separate sheets correspond to each interacting target cell type. Columns include the interaction ID, source and target cell types in control and CK2α knockout samples, ligand and receptor identity, differential expression statistics, ligand and receptor complex status, mean expression and proportion of expressing cells, ligand-receptor interaction scores, CellPhoneDB significance values, interaction weights, specificity and magnitude rankings, and significance of each interaction in control and knockout conditions.

**Table S8. Ligand-receptor interactions with bone marrow HSCs as target:** This table contains predicted ligand-receptor interactions in which bone marrow HSCs act as the target cell type. Separate sheets correspond to each interacting source cell type. Columns include the interaction ID, source and target cell types in control and CK2α knockout samples, ligand and receptor identity, differential expression statistics, ligand and receptor complex status, mean expression and proportion of expressing cells, ligand-receptor interaction scores, CellPhoneDB significance values, interaction weights, specificity and magnitude rankings, and significance of each interaction in control and knockout conditions.

**Table S9. Ligand-receptor interactions with spleen HSCs as source:** This table contains predicted ligand-receptor interactions in which spleen HSCs act as the source cell type. Separate sheets correspond to each interacting target cell type. Columns include the interaction ID, source and target cell types in control and CK2α knockout samples, ligand and receptor identity, differential expression statistics, ligand and receptor complex status, mean expression and proportion of expressing cells, ligand-receptor interaction scores, CellPhoneDB significance values, interaction weights, specificity and magnitude rankings, and significance of each interaction in control and knockout conditions.

**Table S10: Ligand-receptor interactions with spleen HSCs as target:** This table contains predicted ligand-receptor interactions in which spleen HSCs act as the target cell type. Separate sheets correspond to each interacting source cell type. Columns include the interaction ID, source and target cell types in control and CK2α knockout samples, ligand and receptor identity, differential expression statistics, ligand and receptor complex status, mean expression and proportion of expressing cells, ligand-receptor interaction scores, CellPhoneDB significance values, interaction weights, specificity and magnitude rankings, and significance of each interaction in control and knockout conditions.
